# Development and optimization of expected cross value for mate selection problems

**DOI:** 10.1101/2024.05.26.595981

**Authors:** Pouya Ahadi, Balabhaskar Balasundaram, Juan S. Borrero, Charles Chen

## Abstract

In this study, we address the mate selection problem in the hybridization stage of a breeding pipeline, which constitutes the multi-objective breeding goal key to the performance of a variety development program. The solution framework we formulate seeks to ensure that individuals with the most desirable genomic characteristics are selected to cross in order to maximize the likelihood of the inheritance of desirable genetic materials to the progeny. Unlike approaches that use phenotypic values for parental selection and evaluate individuals separately, we use a criterion that relies on the genetic architecture of traits and evaluates combinations of genomic information of the pairs of individuals. We introduce the *expected cross value* (ECV) criterion that measures the expected number of desirable alleles for gametes produced by pairs of individuals sampled from a population of potential parents. We use the ECV criterion to develop an integer linear programming formulation for the parental selection problem. The formulation is capable of controlling the inbreeding level between selected mates. We evaluate the approach for two applications: (i) improving multiple target traits simultaneously, and (ii) finding a multi-parental solution to design crossing blocks. We evaluate the performance of the ECV criterion using a simulation study. Finally, we discuss how the ECV criterion and the proposed integer linear programming techniques can be applied to improve breeding efficiency while maintaining genetic diversity in a breeding program.

## INTRODUCTION

Plant and animal breeding consists of methodologies for the creation, selection, and fixation of superior phenotypes to fulfill the breeding goals of increasing productivity and financial returns, improving welfare, and reducing environmental impact Oldenbroek and van der Waaij (2015). Traditionally, breeders achieve these goals by identifying the individuals with desirable phenotypes and crossing them to produce the segregation of phenotypes in a new generation that allows further selection for advancement. This breeding strategy is perpetuated because high-volume crossing and evaluation led to the identification of the iconic *Green Revolution* varieties that successfully doubled rice and wheat yields from the 1960s to 1990s (Hesser 2006), despite the inevitable inefficiency of producing a high number of failed crosses. However, the future of food security and livestock will be driven not only by the demand but also by severe competition with other uses of land and water resource (Cassandro 2020). Therefore, more efficient breeding strategies ought to be considered because making many crosses with the knowledge that most fail is not justified either by theory or comparative experiments, and is also socially unacceptable.

Ultimately, the overall objective of a breeding program is to produce lines and varieties that are genetically homogeneous and perform at a high level, with end-use quality supportive of the intended market class. A wheat breeding pipeline, for instance, would begin with assembling parental stocks with a careful examination of available germplasm and donor traits. In principle, this is to construct and partition parental stocks respective to a specific goal or goals, to create the genetic variability needed for producing an adapted, high-yielding pure-line variety with perceived quality demands in the future marketplace.

With the continued advancement of genomic technologies and steady decline in genotyping costs, breeders are now able to take full advantage of the availability of genetic information embedded in the genome (Heffner et al. 2010; Hayes et al. 2009). Nevertheless, except for the potential application of a higher selection intensity with GEBVs (genomic estimated breeding values) (Meuwissen et al. 2001), experimental data for the optimal number of crosses as well as the optimal numbers of progeny to sample from each cross required for selection as the initial investment to fulfill the breeding goal have not been reported in the literature (Donald 1968). This is unsurprising given that the number of individuals a breeding program can phenotypically evaluate is resource-limited (Rincent et al. 2017). For example, consider a single cross of two genetically distinct parental lines with 100 QTLs associated with variability among multiple desirable traits. Assuming independent assortment and co-dominance, the complete population from this single pair of founders will consist of 3^100^ ≈ 5.1 × 10^47^ genotypic combinations. Even considering a moderate number of 200 wheat lines in the parental stocks, the number of combinations to be evaluated in the field is astronomically high (Beans 2020). Consequently, analytical approaches based on operations research, mathematical optimization, and statistical learning to optimize breeding decision-making have gained prominence over the years (Johnson et al. 1988; Byrum 2015; Byrum et al. 2016, 2017; Kusmec et al. 2021).

There are two essential steps to addressing this problem using mathematical optimization. The first is to define a fitness criterion to evaluate individuals or crosses based on genetic information. The second step is to devise a mathematical optimization framework that incorporates the fitness criterion along with other essential requirements of the breeding program, and whose objective is to find the individuals or crosses that maximize the fitness criterion. The mathematical optimization framework, while faithfully capturing the various breeding requirements and objectives, must also be computationally viable in order for it to be useful in practice.

In contrast to addressing these breeding challenges in a traditional phenotype-centric paradigm, genetic improvement can also be more efficiently achieved by transferring desirable alleles from parents to progeny as a genetic process while avoiding alleles that show antagonistic pleiotropic effects. Therefore, our aim is to devise a multi-objective mathematical optimization framework that targets more than one phenotype and generates multiple crosses that identify a set of best parental pairs from populations to address multiple breeding goals simultaneously. As it is to be expected in any non-trivial multi-criteria decision-making setting, the criteria (breeding objectives) can be mutually conflicting, making it challenging to design an effective multi-objective optimization framework. For example, yield production in wheat has been found to be negatively correlated with grain protein content (Simmonds 1995), which is an essential factor for its commercial demand (Visscher et al. 1996). This makes concurrently fulfilling breeding goals of high yielding and protein content difficult. The negative correlation between the mass of beef cows and various measures of fertility and stayability could have attributed to the increasing concerns about compromised reproductive efficiency as a result of selection for growth (Berry and Evans 2014; Mwansa et al. 2002).

In this study, we propose a new fitness criterion called the *expected cross value* (ECV) that is inspired by a related fitness criterion called *predicted cross value* (PCV) introduced by Han et al. (2017). ECV returns a probabilistic measure of the fitness of the progeny of a specific pair of individuals based on the genetic architecture of trait variation. We consider the complexity of genetic architecture that underlies agronomic performance characteristics and develop an integer linear programming formulation of the parental pair selection problem that optimizes the ECV criterion. We further extend its capability to select multiple pairs of parents. Our optimization framework is based on the genetic transmission of all detectable genetic loci and can mitigate the potential impact of crossing within highly related individuals. Based on simulation studies, we demonstrate that using ECV as a fitness criterion would address the limitations of other related approaches for mate selection problems, and our multi-objective methodology can simultaneously improve a group of target phenotypic traits.

## METHODS

We begin with some preliminaries needed to formally define the *expected cross value* (ECV) as a new fitness criterion for the mate selection problem. Considering all diploid and polyploid species that may behave as diploids cytologically, e.g., bread wheat (Riley and Chapman 1958), we assume that the variability of target traits is governed by segregating alleles at *N* different loci of all chromosomes in the genome. We use the index set notation [*a*]: = {1, 2, …, *a*} for a positive integer *a*, and define the genotype matrix next.

### Definition 1.

*Given an individual k, we define its genotype matrix L*^*k*^ *as an N* × 2 *binary matrix with the i*, *j-th entry for every i* ∈ [*N*] *and j* ∈ [2] *given by:*

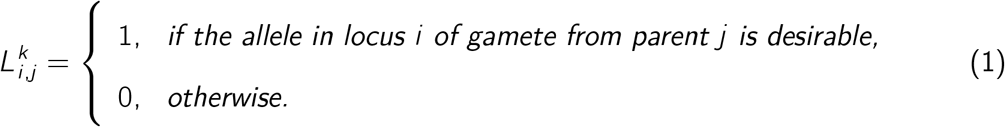

Genotype matrix information of all individuals is an input for the ECV. Hereafter, we refer to the allele in QTL *i* as the *i* -th allele for ease of discussion. In our simulations, alleles are desirable when they enhance the trait value, assuming larger the positive value, the better. Observe that each column of *L*^*k*^ represents a gamete from one of the parents of individual *k*.

We model how alleles transfer from parents to children, i.e., how a gamete inherits alleles from the parent, by using a random *N*-dimensional binary vector *J*, with each component being a random variable *J*_*i*_ for each *i* ∈ [*N*] defined as follows:

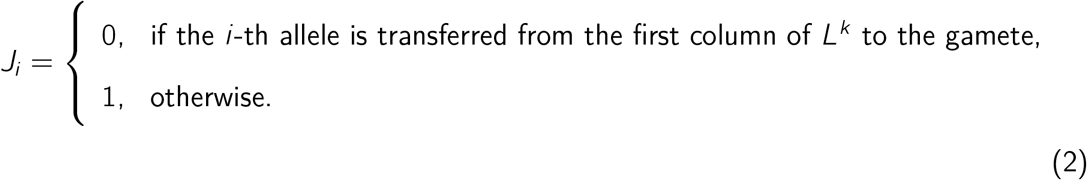

For a given individual *k* and QTL *i*, random variable *J*_*i*_ determines which of the gametes that comprise the genome of individual *k* will transfer the allele in the *i* -th locus.

### Definition 2

(Han et al. (2017)). *We say that the random vector J* ∈ {0, 1}^*N*^ *follows an inheritance distribution with parameters r* ∈ [0, 0.5]^*N−*1^ *and α*_0_, *denoted by J* ∼ ℐ(*r, α*_0_), *if and only if*

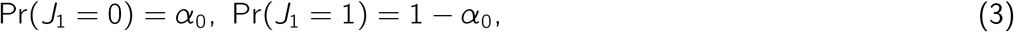

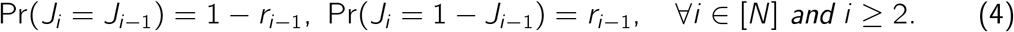

In Definition 2, the (*N* −1)-dimensional vector *r* ∈ [0, 0.5]^*N−*1^ represents the recombination frequencies between the consecutive pairs of loci. The value of *r*_*i*_ is the probability that the *i* -th and (*i* + 1)-th alleles come from different gametes that comprise the genome of individual *k*. Note that if *r*_*i*_ = 0 for all *i* ∈ [*N* − 1], then the gamete produced by individual *k* is identical to exactly one of the parental gametes, while the maximum possible recombination between gametes is expected to be observed when *r*_*i*_ = 0.5 for all *i* ∈ [*N* − 1].

### Deriving the closed-form marginal inheritance distributions

Given *J* ∼ ℐ(*r, α*_0_), we now derive the marginal distribution of *J*_*i*_ for each *i* ∈ [*N*]. The closed-form expressions so obtained then allow us to compute the expectations required to obtain a general closed-form expression for the ECV. For each *i* ∈ [*N*], define the recursive function *φ*_*i*_: ℝ^*N−*1^ − → ℝ as follows:

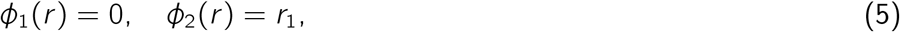

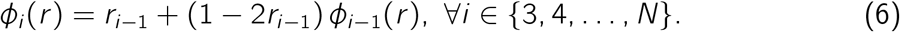

#### Proposition 1.

*Suppose that J* ∼ ℐ(*r, α*_0_). *Then, for each i* ∈ [*N*], *the marginal distribution of J*_*i*_ *satisfies the following equations:*

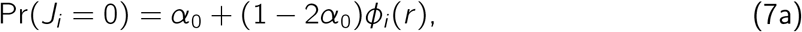

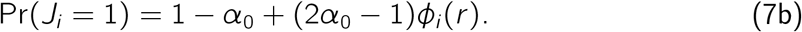

Proposition 1 (proved in the Supplement) establishes the marginal distributions through a recursion, which can be used to obtain a closed-form expression. This result can be further simplified using the laws of inheritance that the allele pairs of a locus segregate randomly during meiosis, and each allele transmits to the gamete with equal probability. Specifically, Proposition 1 then implies the following corollary.

#### Corollary 1.

*Assume that Mendel’s second law holds and α*_0_ = 0.5. *Then*,

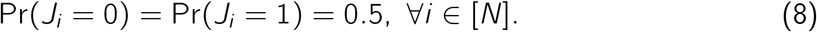

*Furthermore*,

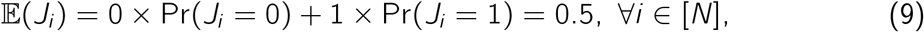

*where 𝔼*(·) *represents the expectation operator*.

### The gamete and loss functions

The inheritance distribution characterizes the source of alleles transmitted from a parent to its gametes. Therefore, we can define a so-called *gamete function* to specify the alleles in the gamete according to the inheritance distribution. Given this gamete function, we derive a closed-form expression for the ECV of a pair of individuals.

#### Definition 3

(Han et al. (2017)). *Given an individual with genotype matrix L and a vector J* ∼ ℐ(*r, α*_0_), *the vector-valued gamete function* gam: (*L, J*) 1→ *g outputs the binary gamete vector g defined as follows for each i* ∈ [*N*]:

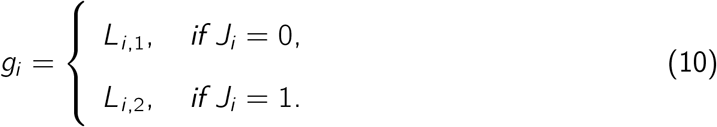

*Equivalently, g*_*i*_ = *L*_*i*,1_(1 − *J*_*i*_) + *L*_*i*,2_*J*_*i*_ .

Suppose we have two individuals with genotype matrices *L*^1^ and *L*^2^, and two independent random vectors *J*^1^, *J*^2^ ∼ ℐ(*r, α*_0_). By crossing these two individuals, the genotype matrix for a child in the progeny is given by matrix [*g*^1^, *g*^2^] where *g*^1^ = gam(*L*^1^, *J*^1^) and *g*^2^ = gam(*L*^2^, *J*^2^). Then, the gamete that is produced by a child of this progeny for the next generation is given by:

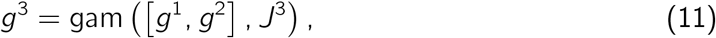

where *J*^3^ ∼ ℐ(*r, α*_0_) is independent of *J*^1^ and *J*^2^. Below, we define a *loss function* in terms of the *g*^3^ gamete vector that will lead us to the ECV criterion.

#### Definition 4.

*Suppose L*^1^ *and L*^2^ *are the genotype matrices of two individuals and let J*^*k*^, *k* = 1, 2, 3, *be independent random vectors following the distribution ℐ*(*r, α*_0_) *for some given r and α*_0_. *Let g*^*k*^ = gam(*L*^*k*^, *J*^*k*^) *for k* = 1, 2 *and g*^3^ = gam([*g*^1^, *g*^2^], *J*^3^). *We define the loss function associated with L*^1^, *L*^2^, *r*, *and α*_0_ *as the following random variable:*

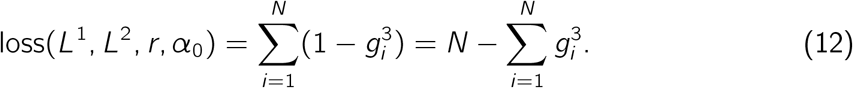

The loss function counts the number of undesirable alleles in the gamete *g*^3^. If the loss function is equal to 0, then all alleles in *g*^3^ are desirable, while the opposite is true if it is equal to *N*. Before deriving our ECV criterion, we introduce the related PCV criterion of Han et al. (2017).

#### Definition 5

(Han et al. (2017)). *Let L*^1^ *and L*^2^ *be the genotype matrices of two individuals, and let r and α*_0_ *be given. Define the gamete g*^3^ *using Equation* (11). *Then, the PCV associated with L*^1^, *L*^2^, *r*, *and α*_0_ *is the probability that the gamete g*^3^ *contains only desirable alleles. That is*,

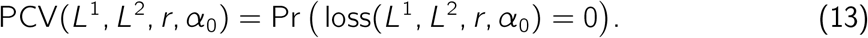

### The expected cross value criterion

Next, we use the loss function in Definition 4 to define the ECV, an alternative criterion to PCV, based on allelic information of individuals. The measure depends on the gamete *g*^3^ defined in Equation (11) and can evaluate a pair of individuals that could be mated.

#### Definition 6.

*For a selected pair of individuals with genotype matrices L*^1^ *and L*^2^, *the ECV is the expected number of desirable alleles in gamete g*^3^ *defined in Equation* (11). *As the loss function represents the number of undesirable alleles in g*^3^, *the ECV can be computed as:*

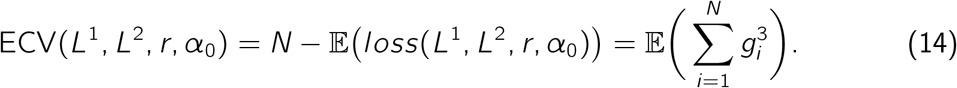

A pair of individuals with the highest ECV value could be selected as parents for crossing. Theorem 1 (proved in the supplement) constitutes our main result that provides a closed-form expression for calculating ECV for a pair of parents.

#### Theorem 1.

*Assume Mendel’s second law holds true and let L*^1^ *and L*^2^ *be the genotype matrices of two individuals. The ECV corresponding to the desired phenotypic trait can be computed using the following equation:*

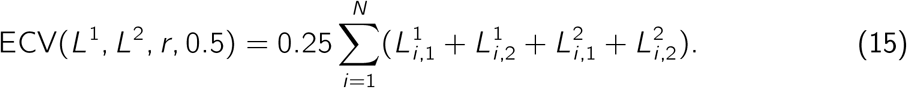

#### Remark 1.

*Without relying on Mendel’s second law, the ECV can still be computed in closed-form more generally as:*

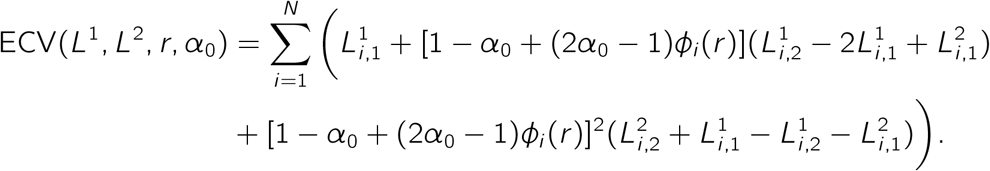

Theorem 1 provides a closed-form expression for the ECV criterion that enables us to formulate the parental selection problem as an integer linear program.

### Single-trait parental selection problem

We develop an integer programming (IP) formulation for the parental selection problem using the ECV criterion as the single optimization objective (see Supplementary Formulation (27)) and the constraint system (and decision variables) from the mixed-integer programming formulation for the PCV introduced by Han et al. (2017). The formulation finds the best pair of individuals maximizing the ECV criterion based on a desired phenotypic trait. In addition, we restrict the inbreeding between selected individuals by preventing pairs of individuals with a sufficiently large inbreeding value from being selected as parents. By using the marker genotype information we can construct the genomic matrix *G* that quantifies the genomic relationship between any pair of individuals in the population (VanRaden 2008). Any pair in the population that has a genomic relationship (i.e., inbreeding value) higher than a pre-determined parameter *ϵ*, will be excluded from the set of feasible pairs using a family of constraints we include in the formulation.

In a breeding program, we may also seek to find multiple pairs for crossing, rather than just a single pair. In order to do so, we introduce Supplementary Algorithm 1 that iteratively solves our IP formulation for the single-trait parental pair selection problem. Note that solving the Supplementary Formulation (27) will identify a pair of individuals as the optimal solution for the problem. By adding “conflict constraints” corresponding to this optimal pair to the formulation, we can exclude *just* this optimal solution from the set of feasible solutions and reoptimize to find the next optimal pair. We can repeat this process until the required number of pairs have been chosen for the crossing program (assuming that many solutions exist).

The flowchart in Figure 1 illustrates the workflow of the proposed ECV approach for mate selection problems for a single trait. The process begins with an initial population where genetic marker and QTL information are available for the selection of parental lines to assemble the crossing block to advance specific breeding targets (Velu and Singh 2013). The ECV criteria can be optimized over several generations (denoted by *T* in Figure 1). In each generation, genomic information related to QTLs and genetic markers, along with a genetic relationship matrix (*G*) is used for constructing the optimization model detailed in Supplementary Formulation (27), and solving it to find an optimal set of mating pairs for crossing. The workflow for solving the multi-trait parental selection by optimizing the ECV metric mirrors the process in in Figure 1 for single-trait ECV optimization. The key difference is that we solve the Supplementary Formulation (29) via lexicographic optimization with user-specified degradation tolerances as described in detail in the Supplement.

**Figure 1:**
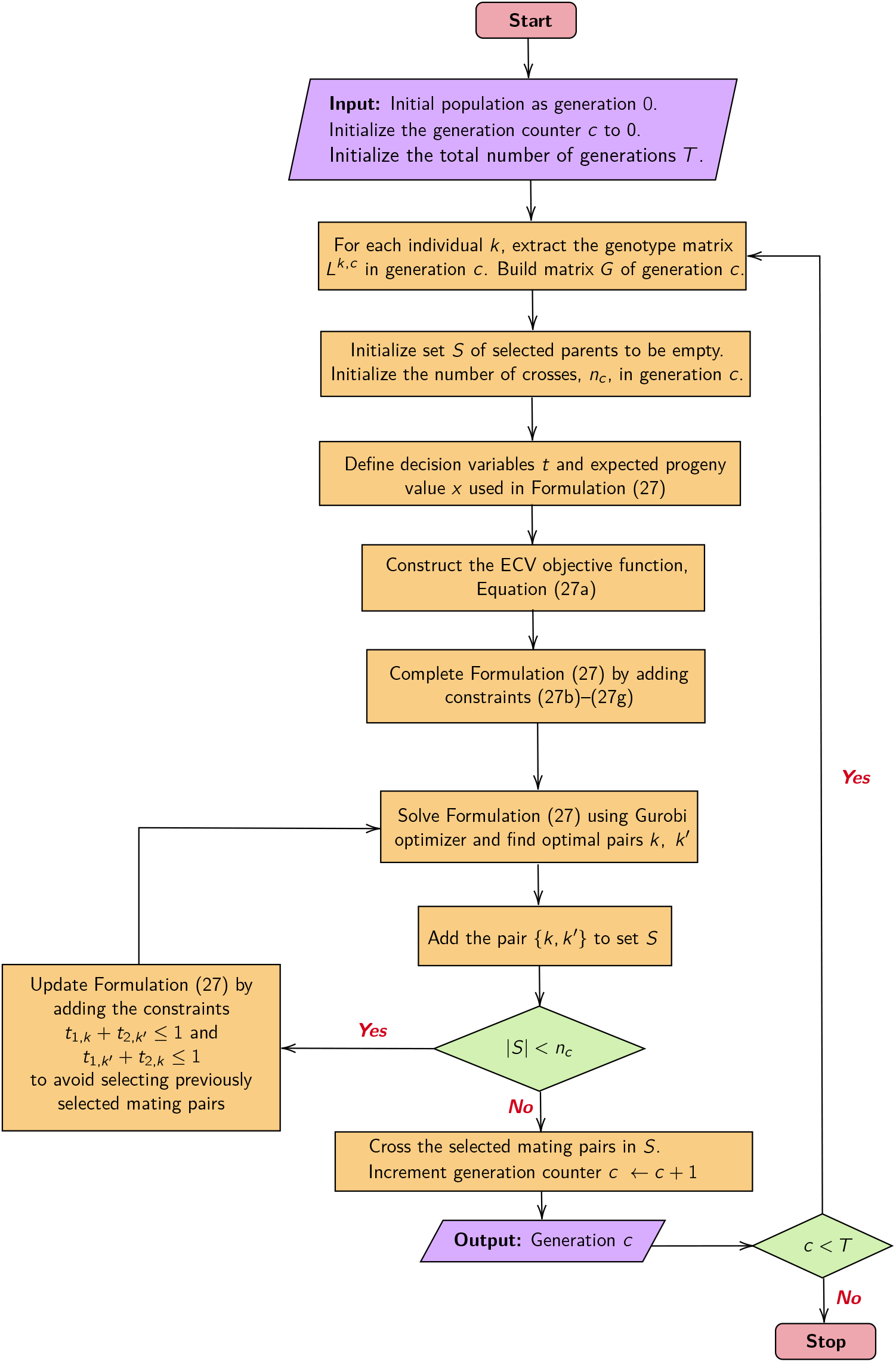
A flowchart describing the overall process for single-trait ECV optimization producing *n*_*c*_ mating pairs in each generation *c* = 1, 2, …, *T*. The optimization formulation and equations referred in the flowchart are included in the Supplementary Information.

### Multi-trait parental selection problem

In general, breeders may be interested in improving several phenotypic traits simultaneously. In this case, we need to extend the ECV criterion to account for multiple traits. We assume there are *M* target traits in the breeding program and that the *ℓ*-th desired trait for every *ℓ* ∈ [*M*] is affected by *N*_*ℓ*_ different QTL in the genome. For each individual we define *M* genotype matrices, one for each trait. Each such matrix is an *N*_*ℓ*_ × 2 binary matrix in which each row represents the pair of alleles in the corresponding genetic locus. Thus, we extend the previous definitions as follows.

#### Definition 7.

*For k* ∈ [*K*] *and ℓ* ∈ [*M*], *the genotype matrix L*^*k*,*ℓ*^ *associated with the k-th individual and the ℓ-th trait is defined as:*

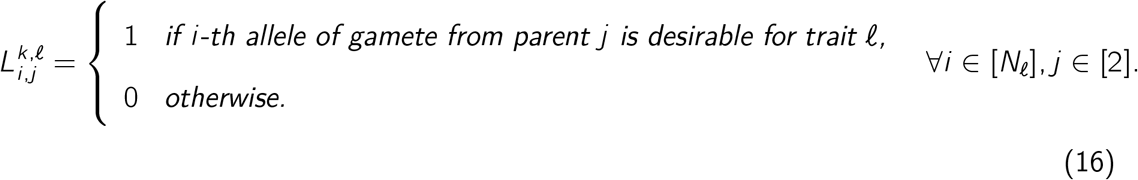

Consider two individuals with genotype matrices *L*^1,*ℓ*^ and *L*^2,*ℓ*^ for target trait *ℓ* ∈ [*M*], and suppose that we have three independent random vectors *J*^1^, *J*^2^ and *J*^3^ following an inheritance distribution ℐ(*r, α*_0_). Using the definition of gamete function (10), the genotype matrix corresponding to the *ℓ*-th trait for a child in the progeny is represented by matrix [*g*^1,*ℓ*^,*g*^2,*ℓ*^] where *g*^1,*ℓ*^ = gam(*L*^1,*ℓ*^, *J*^1^) and *g*^2,*ℓ*^ = gam(*L*^2,*ℓ*^, *J*^2^). The gamete that is produced by this progeny for the next generation is then given by:

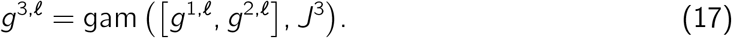

#### Definition 8.

*For the ℓ-th target trait and a selected pair of individuals with genotype matrices L*^1,*ℓ*^ *and L*^2,*ℓ*^, *the ECV* ^*ℓ*^ *is the expected number of desirable alleles of trait ℓ in gamete g*^3,*ℓ*^. *Following Equation* (17), *ECV* ^*ℓ*^, *for each ℓ* ∈ [*M*] *is defined as:*

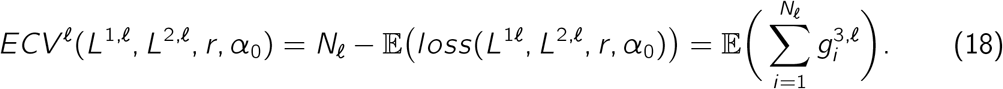

Following Theorem 1, we can obtain a closed-form expression for *ECV* ^*ℓ*^ function.

#### Theorem 2.

*Assume Mendel’s second law holds true and let ℓ* ∈ [*M*]. *Then, for a selected pair of individuals with genotype matrices L*^1,*ℓ*^ *and L*^2,*ℓ*^, *the ECV corresponding to the ℓ-th target phenotypic trait can be computed as:*

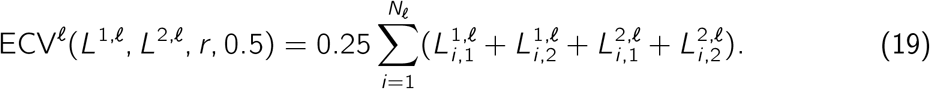

Ideally, a breeding program would like to select parental pair(s) that simultaneously optimize all the ECV^*ℓ*^ functions. Such an optimum is not likely to exist in practice because some phenotypic traits are negatively correlated. Therefore, improving one trait might worsen others. In order to achieve a reasonable trade-off, one turns to the theory of multi-objective optimization.

Consider a vector of objective functions *F* (*t, x*) = ⟨*f*_1_(*t, x* ^1^), *f*_2_(*t, x* ^2^), …, *f*_*M*_(*t, x* ^*M*^)⟩ where *f*_*ℓ*_(*t, x* ^*ℓ*^) denotes the ECV function (19) corresponding to *ℓ*-th trait. Supplementary Formulation (29) for the multi-trait parental selection problem seeks to find a pair of individuals that will “maximize” the vector-valued objective function. Similar to the single-trait optimization model, this formulation also excludes pairs of individuals with genomic relationship exceeding the tolerance threshold from the set of feasible solutions. Furthermore, as explained in the previous section, this approach can also be extended to select multiple parental pairs for the breeding program by iteratively adding conflict constraints. The differences lie in the handling of multiple traits, especially in the vector objective function.

Multi-objective or vector optimization problems are commonly handled by scalarization— converting the vector optimization problem into one or more scalar optimization problems (Miettinen 2012; Sawaragi et al. 1985); see survey by Miettinen et al. (2016) for interactive and other methods. One approach is to use a weighted combination of the individual objective functions to produce an optimization problem with a scalar objective. The weights, which are predetermined by the user, need to be carefully chosen to ensure they reflect the relative importance of the individual objectives and also scale them appropriately as necessary. Another approach, *lexicographic optimization*, prioritizes the objective functions based on their importance and optimizes them sequentially, starting with the most important. While optimizing lower priority objectives, we restrict the feasible region to only those solutions that will not degrade the higher priority objectives, or limit their degradation by user-specified tolerances.

The weighted sum approach, where we aggregate the individual objectives into a single objective using user-defined weights, requires a vector of weights that capture the importance of each phenotypic trait in the breeding program. In practice, it is difficult to identify a precise and meaningful weight for each trait as there are many factors of the breeding program (some of them potentially unknown) that might play a role in defining it. By contrast, it might be simpler for a breeding program to order the traits based on their importance.

The lexicographic optimization approach is not without drawbacks, as it could degenerate into single-objective optimization with the highest priority objective if we subsequently allow no degradation of higher priority objectives. In the worst case, if the first objective has a unique optimal solution and we tolerate no degradation on the first objective, the subsequent objectives are irrelevant. The use of tolerance is therefore important as it allows limited degradation of a higher priority objective when optimizing a lower priority objective, but allows for a larger feasible solution space for the lower priority objective (when compared against using zero tolerance). Hence, we will be using lexicographic maximization with positive tolerances in solving Supplementary Formulation (29).

Assume without loss of generality that the vector of objective functions, 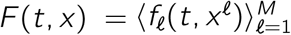, is already in decreasing order of importance. Thus trait *ℓ* is more important than trait *ℓ* + 1, for each *ℓ* ∈ [*M* − 1]. The solver we use in our computational studies is capable of lexicographic optimization with degradation tolerances for objectives specified by the decision maker. Let us denote these tolerances by *τ* = (*τ*_1_, *τ*_2_, …, *τ*_*M*_), where *τ*_*ℓ*_ ∈ [0, 1] for each trait *ℓ* ∈ [*M*]. The solver optimizes the first objective function *f*_1_(*t, x*) and then, among those feasible solutions within a factor (1 − *τ*_1_) of the optimal objective value of the first objective function, optimizes the second objective function. This process is repeated until the last objective is optimized. In particular, this method assures that the optimal solution for the *ℓ*-th objective, for *ℓ* = 2, …, *M*, is within a factor (1 − *τ*_*i*_) of the optimal value of the *i* -th objective, for every *i* ∈ [*ℓ* − 1]. As *f*_*M*_(*t, x* ^*M*^) is the last objective function to optimize, there is no need for a tolerance *τ*_*M*_, and hence we set *τ*_*M*_ = 0 for all our experiments.

### Simulation study

Simulations were conducted to evaluate the performance of ECV compared to other parent selection approaches using phenotypes and breeding values (GEBV). Two simulation experiments were considered in this study. First, we considered a single-trait optimization problem to solely improve Trait 1, simulated as a mixture of traits with oligogenic and polygenic genetic architectures. Next, we examined a multiple-trait parent selection problem where the breeding program was tasked to simultaneously improve all traits of interest. In this experiment, we simulated a polygenic architecture for Trait 3, representing a trait such as yielding capacity that is usually governed by a large number of loci where each allele has a small impact on the expression of the trait and in a negative genetic correlation with Trait 1, in addition to an oligogenic phenotype (Trait 2) that may imitate the genetic architecture underlying disease resistance.

For all experiments, two metrics were reported from the simulations, average desirable allele frequency and average phenotypic trait values of the progeny, to compare the performance of the methods in each generation. We also recorded the average genomic relationship for the selected individuals for all three approaches. In the case of multi-parental pair selection, we sorted pairs of individuals based on the summation of their trait values or GEBVs and made selection decisions based on the summations of trait values. Moreover, by default, there was no control over the genomic relationship between selected parent pairs for the phenotypic selection and GEBV selection approaches; however, we assumed that self-crossing is not a feasible choice in these approaches.

The QU-GENE engine and QuLinePlus proposed by Ali et al. (2020) were used to simulate initial populations and the progeny in the subsequent generations. The QU-GENE engine establishes the initial population with inputs of genetic effects for segregating alleles, recombination frequencies and the number of desired individuals. We considered an initial population such that the allele frequency at all loci was set at 0.5. In our experiments, QuLinePlus took the genotypic information of a population and a list of selected pairs, simulated the progeny by crossing the selected parental pairs, and output genotypic and phenotypic information for all individuals in the subsequent generation. The GEBVs were calculated using the “rrBLUP” package (Endelman 2011). The Gurobi Optimization Solver (Gurobi Optimization, LLC 2024) was used to solve the integer linear programming formulations that were implemented in the Python programming language.

The initial population consisted of 10,000 individuals, with 200 biallelic genetic loci and 100 markers. Of these, 40 genetic loci had effects on Trait 1, 10 on Trait 2, and 70 on Trait 3. The markers had no genetic effects on any of the traits. Trait 3 and Trait 1 share 20 common loci with pleiotropic effects, which resulted in a negative correlation between those phenotypic traits. We conducted all of the experiments for four generations and for each cross we simulated 100 progeny for the next generation. We performed two sets of simulation studies, assuming a consistent growing environment across generations. In the first simulation study, the number of parental pairs that we chose from the initial population, generations one, two, and three, was 50, 10, 5, and 5, respectively. Thus, the population size in the simulation studies for generations one, two, three and four were, respectively, 5,000, 1,000, 500, and 500, respectively.

To further investigate the effectiveness of our methodology, we explored the impact of selection intensity in our second simulation study. Scenario A imposed a higher selection intensity with 50 crosses made from the initial population (generation 0), and 10, 3, and 3, respectively, for generations one, two, and three. For intermediate selection intensity (Scenario B, same as the first simulation study), from the initial population, generations one, two, and three, we chose 50, 10, 5, and 5 parental pairs, respectively. Finally, in Scenario C, representing a case of reduced selection intensity, the numbers of parental pairs selected in generation one was 25, and 5 parental pairs for the generations two and three.

## RESULTS

The single-trait simulation results over five generations are summarized in Figure 2. For all traits considered, ECV significantly increased the proportion of desirable alleles (see Figure 2a) while showing the capacity to regulate the relatedness within the breeding population by avoiding crossing closely related individuals (see Figure 2c). Further, although statistically insignificant, genetic crosses done by phenotypically superior individuals returned the lowest means of the progeny in all traits, compared to genetics-informed approaches, like GEBVs and ECV (see Figure 2b). However, populations generated by ECV provided a greater potential for advancing individuals with larger phenotypic values.

**Figure 2:**
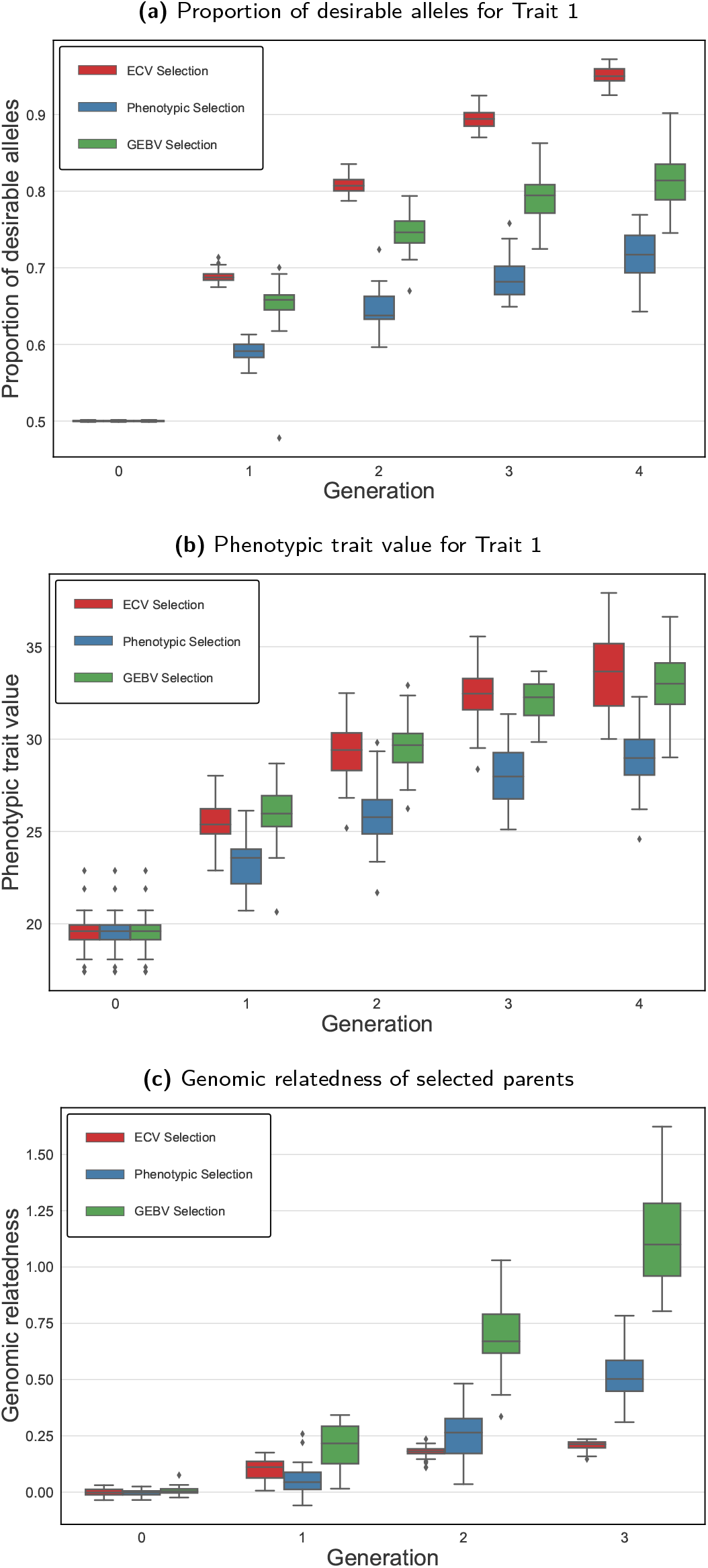
Performance of ECV, phenotypic and GEBV selection methods in 30 simulation runs for single trait improvement.

Single-trait optimization does not guarantee improvement for phenotypes other than the target trait. Figure 3 shows boxplots for Trait 2 and Trait 3 when we optimize Trait 1 in a single-trait ECV optimization framework. The frequency for the desirable allele (Figure 3a) as well as the phenotypic values (Figure 3b) remained unimproved for Trait 2. The scenario could be worse if target traits are determined by QTLs with antagonistic pleiotropic effects. This can be seen in Figures 3c and 3d, which depict a significant decrease in the proportion of desirable alleles and phenotypic values of Trait 3 as a result of optimizing for Trait 1.

**Figure 3:**
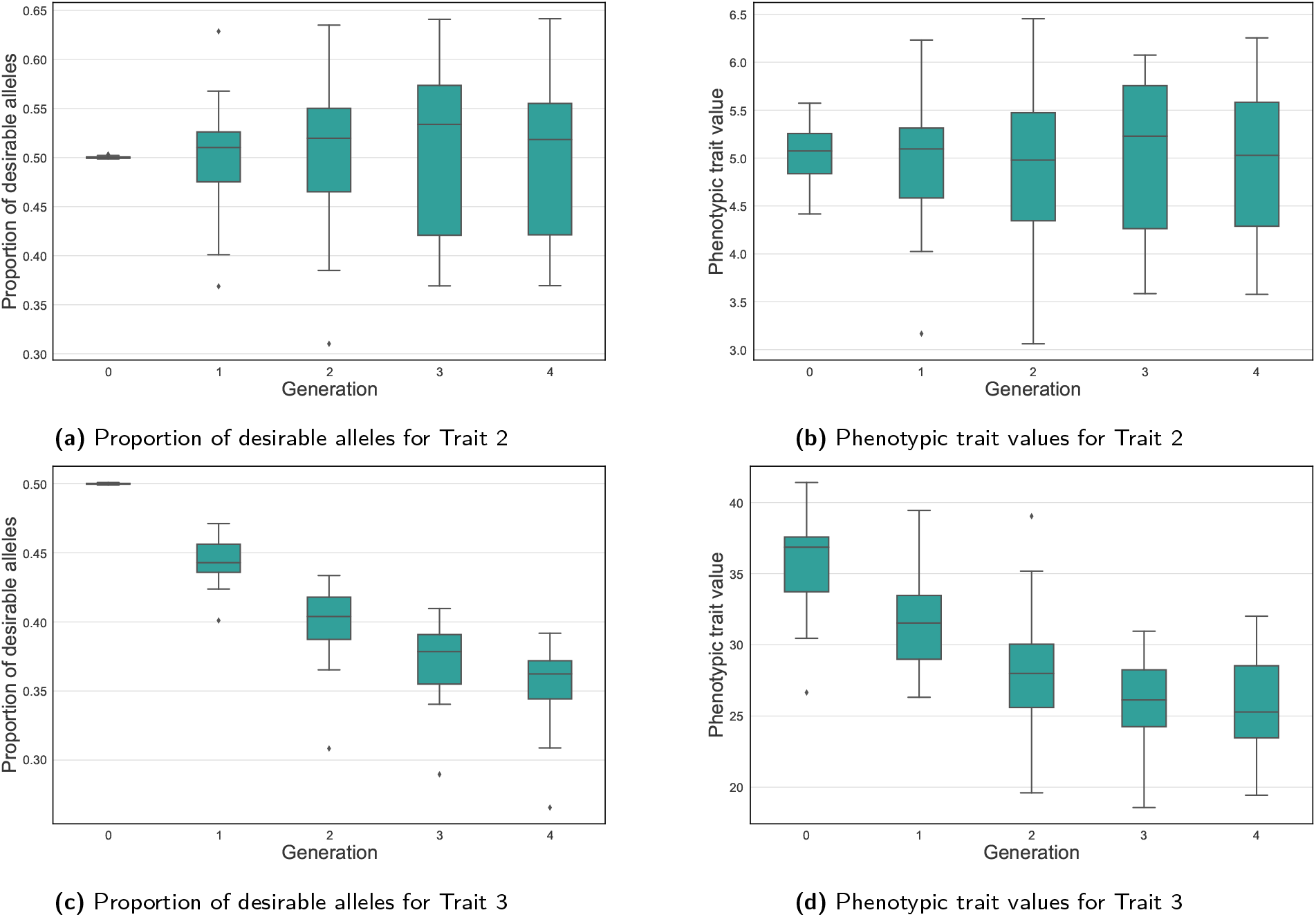
Effect of improving Trait 1 using ECV optimization on the performance of Trait 2 and Trait 3 based on proportion of desirable alleles and phenotypic trait value.

For multi-trait parental selection based on ECV, we employed the lexicographic multi-objective optimization approach described earlier. The tolerances were chosen based on preliminary experiments as follows: let *τ*_*i*,*c*_ denote the degradation tolerance for optimization objective *i* in generation *c*, then we used *τ*_1,0_ = 0.17, *τ*_1,1_ = 0.05, *τ*_1,2_ = 0.05, *τ*_1,3_ = 0.05 and *τ*_2,0_ = 0.00, *τ*_2,1_ = 0.00, *τ*_2,2_ = 0.00, *τ*_2,3_ = 0.05. In general, the tolerance parameters can be calibrated to have the desired impact on the model. The results in Figure 4 show the advantage of using ECV. Despite the negative genetic correlation, ECV was able to increase the desirable allele frequency to 0.70 (±0.02), 0.65 (±0.08), and 0.72 (±0.01), for Trait 1, Trait 2, and Trait 3, respectively. In contrast, the impact of negative correlation between Trait 3 and Trait 1 was most obvious when the phenotypic selection was used, leading to a significant loss of desirable allele for Trait 1 (see Figure 4a). Similarly, ECV improved phenotypic values of the progeny for all traits simultaneously, whereas no improvement for Trait 1 and Trait 2 was found using phenotypic selection in our simulations when the tolerances were set slightly favoring Trait 3. It is noteworthy that a genomics-informed selection method, GEBV, returned comparable results to ECV for Trait 1. This benefit of GEBV, however, is at the expense of genetic diversity, as shown in Figure 5. Genomic relatedness (VanRaden 2008) has increased noticeably over the four generations using the GEBV selection method.

**Figure 4:**
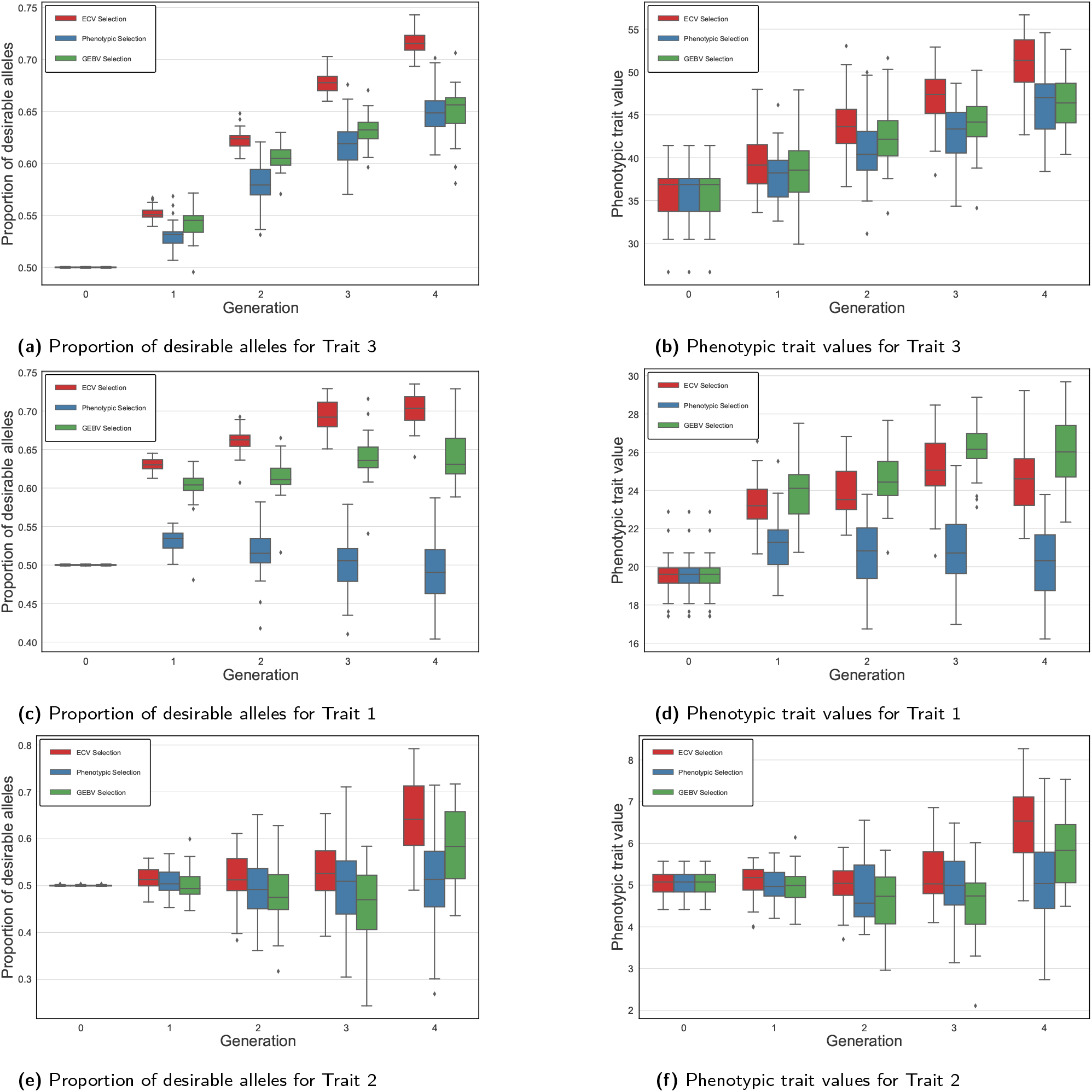
Performance of ECV, phenotypic and GEBV selection methods in 30 simulation runs for multiple trait improvement.

**Figure 5:**
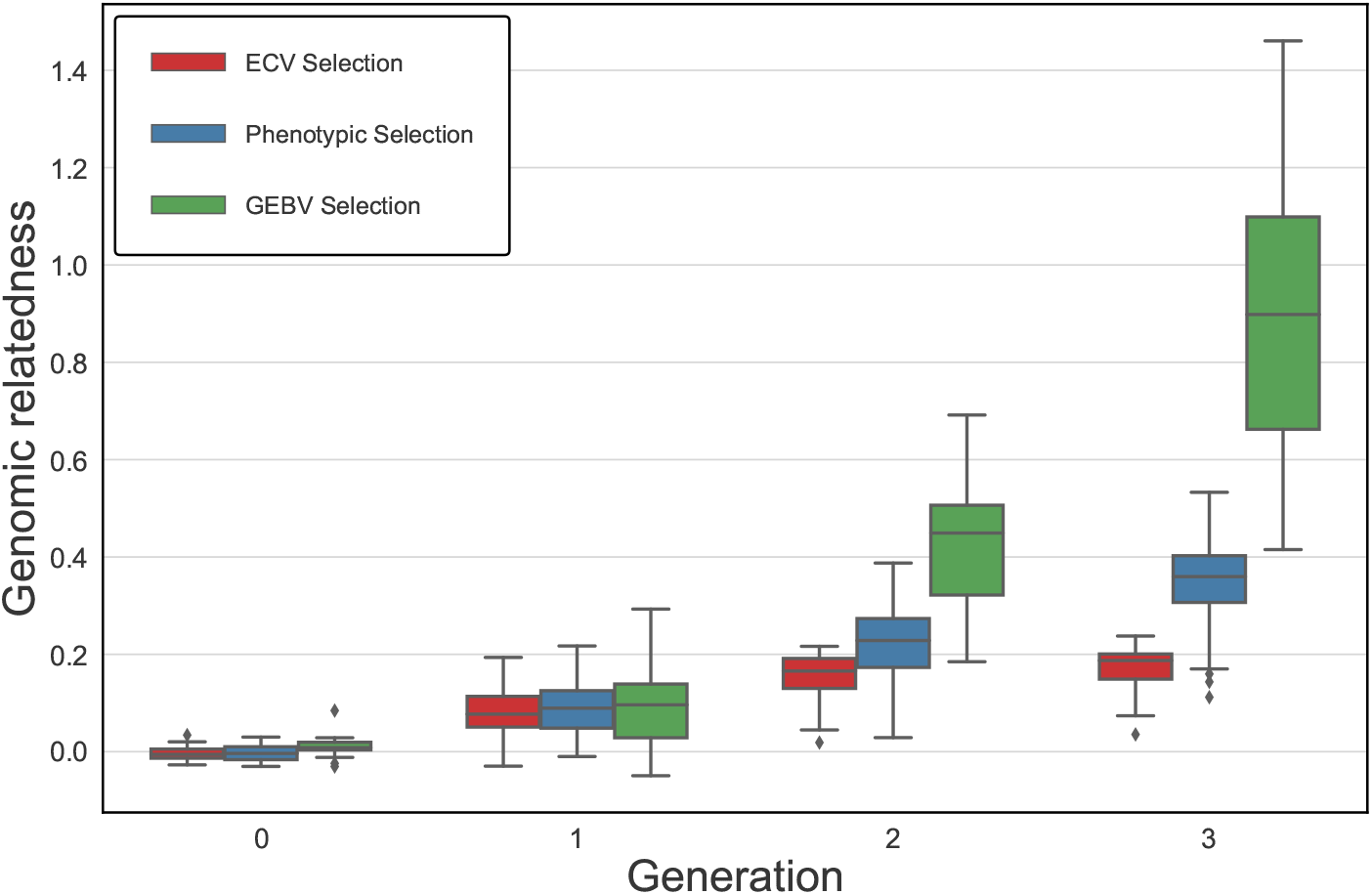
Genomic relatedness for multiple trait improvement based on ECV, phenotypic and GEBV selection methods.

Figure 6 displays the boxplots for the three selection intensity scenarios A, B, and C introduced in the Simulation study, focusing on the proportion of desirable alleles as the performance metric. In the early generations, particularly generation 2, our proposed ECV approach outperformed other selection strategies, most evidently for Traits 1 and 3 in the implementation of multiple trait selection. As the generations advanced, the ECV approach continued to excel in Scenarios B and C, resulting in higher proportion of desirable alleles for Traits 1 and 3. In Scenario A, where selection intensity was higher and ECV selection method was not dominant, the method still yielded replications with a greater proportion of desirable alleles compared to other strategies, despite the genomic relationship constraints inherent in the ECV method. Furthermore, in the last generation under scenario A, the mean (± standard deviation) of genomic relatedness over all replications for the ECV, GEBV, and Phenotypic selection approaches are 0.15 (±0.04), 0.42 (±0.10), and 0.25 (±0.10), respectively. These results illustrate the effectiveness of our ECV optimization framework, with its explicit constraints limiting genomic relatedness, in managing genomic relatedness over generations when selection intensity is higher, while improving desirable breeding traits. The impact of higher selection intensity, however, led to greater variability in the proportion of desirable alleles in Trait 3 of Scenario A, which could be due to the drift effect of the smaller breeding population size (Turner-Hissong et al. 2020).

**Figure 6:**
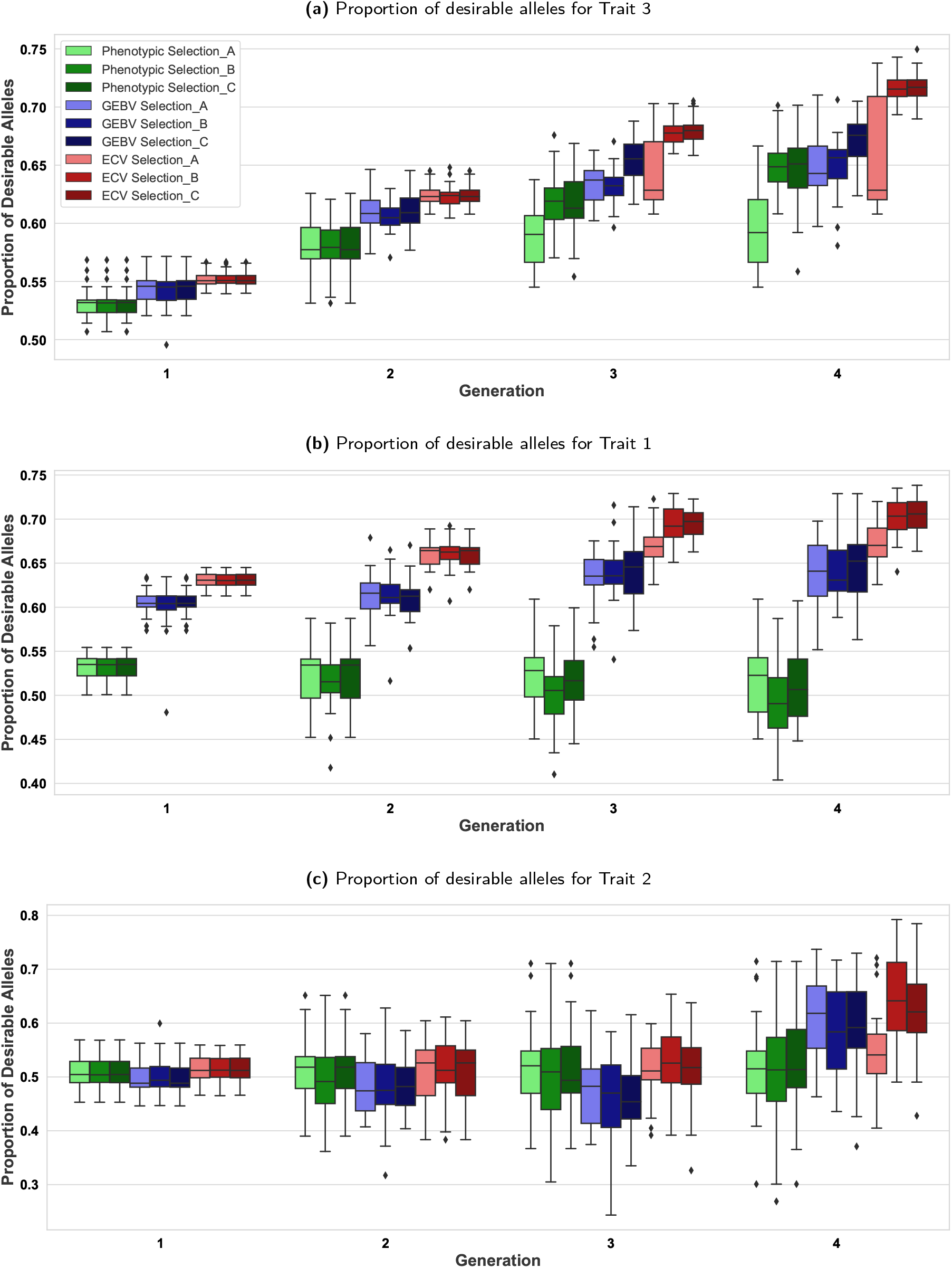
The effect of selection intensity represented by scenario A (higher intensity), scenario B (intermediate intensity), and scenario C (lower intensity) on the proportion of desirable alleles.

## DISCUSSION

The principal objective of breeding is to combine as many desirable traits as possible into a genotype that can be distributed to farmers, producers or breeders. For example, in plant breeding the breeders are tasked with developing elite genotypes that display desirable use characteristics including high yields, disease resistance, and are also well-adapted to a range of environmental conditions (Breseghello and Coelho 2013). These desirable characteristics are typically possessed by multiple founders. By mixing and recombining founder genomes, the distribution of these desirable phenotypes observed in the offspring, owing to the segregation of alleles often distributed throughout the genome, allow breeders to identify superior individuals for subsequent breeding, widespread evaluations, and sales.

However, when these desired characteristics differ in variability, heritability, economic importance, and are correlated with other phenotypes and genotypes, effective mating designs capable of improving multiple traits simultaneously can be challenging to identify (Johnson et al. 1988). This breeding process is also ineffective as breeders tend to make hundreds or thousands of crosses, of which only a few are advanced in the subsequent years (Witcombe et al. 2013). Traditionally, these objectives are achieved by breeding from the “best”—the best being determined by their own phenotypic values (Allard 1999; Akdemir et al. 2019). More advanced techniques, such as pedigree-based (Henderson 1984; Gianola and Fernando 1986), marker-based genetic value predictions (Lande and Thompson 1990; Hospital and Charcosset 1997; Bernardo and Charcosset 2006), and mating designs by genomic information (Akdemir and Sánchez 2016) are also available.

Beginning with the work of Johnson et al. (1988), mathematical programming approaches have facilitated the improvement of genetic traits through the use of mathematical optimization models that aid breeders in making better decisions in selecting mating parents. Toro et al. (1991) solved mate selection problems using linear programming techniques and demonstrated the effectiveness of their approach within multiple ovulation and embryo transfer (MOET) breeding schemes for dairy cattle with the help of simulation studies. Jansen and Wilton (1985) addressed the issue of factorial growth in the number of combinations to cross by formulating and solving an integer programming model to improve the overall progeny merit.

Moeinizade et al. (2019) recently proposed a single-trait optimization of a “look ahead” metric that focuses on a predetermined terminal generation to optimize mating decisions for maximizing expected GEBVs in the terminal generation without explicitly considering the impact of genetic erosion. Amini et al. (2021) further improved this look-ahead framework by prioritizing best individuals for crossing and using multiple prediction algorithms to improve prediction accuracy. These approaches are also complemented by Zhang and Wang (2022) who proposed a “net present value” inspired mechanism for discounting future gains, which values early-term genetic gains more than those anticipated in the future. This was done to overcome a drawback of the original look-ahead scheme by Moeinizade et al. (2019), which can produce slow genetic gains in the early generations and accelerating more rapidly as we approach the terminal generation.

Byrum et al. (2016) report on their long-term development and quantification of an unbiased genetic gain performance metric, and pioneered its use in evaluating breeding projects as varieties were developed. Byrum et al. (2016) and Byrum et al. (2017) demonstrate the successful commercial use of advanced analytics and operations research tools such as integer linear programming, Monte Carlo simulation, and stochastic optimization by the agriculture industry, which has served to further motivate its broader use in many areas of crop and animal sciences; see also (Byrum 2015, 2016).

Furthermore, when the breeding objective involves more than one trait, a selection index of progeny merit was considered as a linear function of estimated breeding values for each trait by Allaire (1980). In animal breeding, for instance, the genetic merit of calves is estimated as half of the sire’s and half of the dam’s breeding value. An optimization-based procedure for mate selection in animal breeding is introduced by Kinghorn (1998, 2011) based on a mate selection criterion proposed by Kinghorn and Shepherd (1999).

In the genomics era, the parental selection problem has been increasingly addressed with the use of genomic relationships (Sun et al. 2013), heuristic searches for gene pyramiding (De Beukelaer et al. 2015), and by modeling the recombination of desirable alleles as a result of crossing (Han et al. 2017; Moeinizade et al. 2019). For the purpose of introgressing a small number of desirable alleles from a donor to a recipient, Han et al. (2017) proposed an efficient algorithm for calculating the PCV defined as the probability that a gamete of a random progeny from crossing two genetic individuals would consist only of desirable alleles. In a specific case where the desirable allele for the *i* -th locus is not present in both parents (denoted by *k* and *k*^*′*^), such that 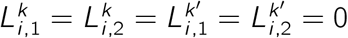, PCV will conclude that the *i* -th component of gamete *g*^3^ is zero with probability one and hence *P CV* (*L*^*k*^, *L*^*k*^*′*, *r, α*_0_) = 0. In this case, the individuals *k* and *k*^*′*^ will not be selected, regardless whether or not there maybe be desirable alleles present in the rest of the genome. While such a result is desirable for the goals of introgressing a small number of alleles for traits like herbicide, disease, or insect resistance, it would be inappropriate for identifying crosses that will have the best opportunity to combine a large number of genetic alleles. Considering the polygenic inheritance of agronomical performance traits (Lynch and Walsh 1998; Scott et al. 2021), the PCV approaches zero for all breeding parents as the number of loci with desirable alleles increases. Consider the following probabilistic inequality (Fréchet inequality):

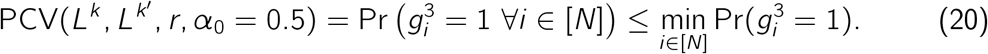

Hence, the larger the value of *N*, the greater the chance 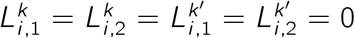 for some QTL *i*. The PCV method could therefore lead to indiscriminate mate selection for traits that have hundreds or thousands of loci with desirable alleles because the PCV value is (nearly) zero for essentially any choice of mates. This observation motivated us to introduce our ECV criterion, especially for breeding targets governed by a large number of genetic loci and for non-introgression projects

As Figure 2a shows, our results demonstrated a significantly greater capacity to increase desirable allele frequency compared to the conventional phenotypic selection and the selection done by the genomics-derived GEBV; and, the benefit of using ECV can be realized in as short as two generations. Moreover, the greater range of trait value distribution presents additional opportunities for breeders to identify the superiors for population advancement (see Figure 2b).

Based on our simulations, we observe that the breeding population has gone from unrelated to essentially full-sibs in three generations of selecting breeding parents based on GEBVs (see Figure 2c). Compared to the phenotypic selection, GEBV selection might have manifested a rapid increase of relatedness by crossing individuals closely related to the training population (Bassi et al. 2016; Forutan et al. 2018). Though GEBV selection might show a capacity to provide short-term genetic gain, selecting breeding parents solely by GEBVs would lead to undesirable consequences such as loss of genetic diversity, further diminishing long-term genetic gain (Jannink 2010; Doekes et al. 2018).

To ensure the capacity to preserve multiple genetic lineages, ECV allows for the selection of more than one pair of individuals, and while self-crossing was not allowed in this study, our method permitted the same individual to be crossed with multiple breeding parents as long as the genomic relationship of the parents was not greater than *ϵ*, a parameter that breeders can use to control how much inbreeding is acceptable.

Fundamental to all variety improvement programs is the identification of the most efficient path to reach breeding objectives (Bernardo 2002; Akdemir et al. 2019). However, breeders are usually tasked with combining a suite of traits in addition to yield and growth components. The negative genetic correlations caused by the non-random association of alleles underlying these breeding objectives impose additional challenges, as selecting based on one trait may adversely impact another (Lynch and Walsh 1998). To simultaneously improve multiple traits, phenotype-based selection indices have been widely considered (Hazel and Lush 1942; Hazel et al. 1994; Villanueva and Woolliams 1997; Jannink et al. 2000; Moeinizade et al. 2020). Selecting breeding parents based on a selection index does not necessarily choose the best genetics to recombine; further, since the selection index applied is a weight assignment of target phenotypes, such decisions could result in the loss of beneficial alleles.

In this study, the proposed ECV framework is based on an allele transmission process. Rather than relying on the phenotypes of breeding parents, ECV identifies the crosses with the highest likelihood of transmitting desirable alleles from pairs of parents to the progeny. In the case where multiple traits need to be considered simultaneously, ECV seeks the optimal combination of alleles for all target phenotypes ordered by their importance, while maintaining a customizable tolerance such that QTLs with antagonistic pleiotropic gene action could remain in consideration before the final breeding recommendation is made. Figure 4 and 6 showed that despite the negative correlation between Traits 1 and 3, ECV was able to increase desirable allele frequency for all traits in our simulation studies. In addition, as seen from Figure 5, the inbreeding coefficient in the progeny was regulated as ECV was optimized with the tolerance constraint on the genomic relatedness between breeding parents. As genotyping has become routine in breeding programs (Hayes and Goddard 2010; Bentley et al. 2022), the application of this constraint ought to be considered to mitigate the multiple trait scenario in Figure 4, where the gain might be built at the expense of genetic diversity (Figure 5), a phenomenon also found in index selection methods (Akdemir et al. 2019). If practical considerations favor breeding parents to be selected from a narrow genetic pool, the constraint could be moved to the objective as a penalty term.

Breeding programs develop elite genotypes that often demonstrate similar essential genomic profiles of desirable end-use characteristics, agronomical attributes, disease resistance packages, as well as adaptation to the target environment. Breeding among the elites can produce new variability as the source of new cultivars with minimal risk of introducing undesirable features. This variation may eventually be exhausted, and new genes and alleles must be introduced. Identifying beneficial alleles from un-adapted material itself has been described as searching for a needle in a haystack (Pixley et al. 2014). Introgressing these novel alleles can also be risky because the unwanted alleles in exotic germplasm may distrupt essential allele combinations (Willcox et al. 2022); and, it requires a higher institutional cost due to a greater number of crosses and longer breeding cycle needed to achieve the breeding objectives (Snelling et al. 2019; Neyhart et al. 2019). Based on our simulations, we reckon that ECV can be an option.

Beyond animal and cereal crop breeding, we suspect that implementing optimization-based methods like ECV could be advantageous to breeding of genetically diverse, long-generation, and slow reaction, cross-pollinated species, such as conifers. Tree breeders generally establish open-pollinated seed orchards for selection (White et al. 2007), and several mating designs have been proposed (Namkoong 1976; Zobel and Talbert 1984), among which the polycross is considered as one of the most cost-effective (Kumar et al. 2007; Lenz et al. 2020). The ability to design the pollen pool while managing inbreeding with ECV will provide the capacity to rapidly increase desirable allele frequencies and, at the same time, avoid severe inbreeding depression for conifer species (Berry and Evans 2014; Mwansa et al. 2002; Snelling et al. 2019).

The conceptual framework we have introduced and our results show that adopting multi-objective optimization tools from operations research to solve breeding problems is highly advantageous (Cameron et al. 2017; Beans 2020; Kusmec et al. 2021). Several improvements or extensions suggested next should also be considered. When the pool of genetic diversity increases, solving integer programming problems for ECV will require further development to account for different distributions of crossover events in different crosses (Stapley et al. 2017; Nachman 2002; Jabbari et al. 2019; Dreissig et al. 2019). Furthermore, as multi-parent populations like MAGIC (multi-parent advanced generation inter-cross) have become a means to provide germplasm for breeding programs (Scott et al. 2020), there is a need to expand optimization frameworks such as ECV to consider multiple parental lineages, which might also help guide the polycross mating design in forestry (Frandsen 1940; Lambeth et al. 2001). Our proposed methodology relies on the the underlying genetic information of the breeding population, such as QTLs and genetic association of desirable traits. While affordable large-scale genotyping and phenotyping technologies are becoming accessible to breeding programs (Reynolds et al. 2020; Bassi et al. 2024), large breeding populations necessitate extensive genomic information, which can be computationally demanding. Moreover, integer linear programming is NP-hard in general, making it challenging to solve very large-scale problems to optimality. In the case of mate selection, the size of the population and the number of genes directly influence the computational time, which implies that massive datasets could make obtaining optimal solutions unrealistic for practical applications. In such circumstances, we may consider modifying our approach to solving the integer linear program by employing decomposition techniques to address the large-scale instances and likely settle for sub-optimal (but good quality) feasible solutions.

Our simulations also indicate that the ECV selection framework may result in higher performance variability when the selection intensifies in earlier generations. To alleviate this issue, we could either relax the genomic relatedness constraint (smaller *ϵ*) in earlier generations or intensify selection only in advanced generations. Further, care must also be taken in choosing the degradation tolerances *τ*_*ℓ*_ for each trait *ℓ* ∈ [*M*] in the lexicographic multi-trait ECV optimization framework, which will entail computational expenditure in terms of preliminary computational experiments, which could become challenging at larger scales.

## ACKNOWLEDGEMENTS

Funding for this work was supported by grants from the Oklahoma Wheat Research Foundation (for CC), Oklahoma Center for the Advancement of Science and Technology (OCAST) award number PS15-011-2 and PS19-004 for CC. This work was completed utilizing the High-Performance Computing Center facilities of Oklahoma State University at Stillwater, and also in part by the Extreme Science and Engineering Discovery Environment (XSEDE), which is supported by National Science Foundation grant number ACI-1548562. Specifically, it used the Bridges system, which is supported by NSF award number ACI-1445606, at the Pittsburgh Supercomputing Center (PSC) under the resource allocation MCB-180177. The authors are grateful to the anonymous reviewers for their careful reading of our original manuscript, their constructive criticism, and providing detailed and thoughtful comments that helped us improve this manuscript.

## AUTHOR CONTRIBUTIONS

BB, JB, and CC were responsible for the conceptualization of the study. PA developed the theoretical results and computer implementations as part of his thesis. PA and CC performed the analysis and wrote the original draft. All authors contributed to interpreting results, providing feedback, and editing and approving the manuscript.

## DATA AVAILABILITY

The data and codes are available at: https://github.com/transgenomicsosu/ECV.

