## supplementary information for "Development and optimization of expected cross value for mate selection problems"

### PROOFS

#### Proof of Proposition 1

We model the random vector $J$ that follows an inheritance distribution as a discrete time Markov chain (DTMC) with $J = \left\{J_n \colon n \geq 0\right\}$ where $J_n$ represents the state of the process at $n$-th step, i.e., the value of the random vector $J$ in the $n$-th position, with the state space $\{0, 1\}$. This process is not a time-homogeneous DTMC. According to Equation (4) in the main article, the transition probability matrix from step $k$ to step $k+1$ is as follows:

$$P_{k:k+1} = \begin{array}{c c} & \begin{array}{cc} 0 & 1 \end{array} \\ \begin{array}{c} 0 \\ 1 \end{array} & \begin{pmatrix} 1 - r_k & r_k \\ r_k & 1 - r_k \end{pmatrix} \end{array} \qquad \forall k \in [N-1].$$

The transition probability matrix from the first step 1 to step $i \in [N-1]$ is then given by:

$$P_{1:i} = \prod_{k=1}^{i-1} P_{k:k+1}.$$

We claim that:

$$P_{1:i} = \begin{bmatrix} 1 - \phi_i(r) & \phi_i(r) \\ \phi_i(r) & 1 - \phi_i(r) \end{bmatrix}, \tag{21}$$

where $\phi_i(r)$ is defined in Equations (5) and (6) in the main article. We prove this claim by induction on $i$. The claim holds for the base case $i = 2$ by definition, because according to Equation (5) in the main article, $\phi_2(r) = r_1$. Let us suppose Equation (21) holds for step

$i = n$. By induction hypothesis, we know that:

$$P_{1:n} = \begin{bmatrix} 1-\phi_n(r) & \phi_n(r) \\ \phi_n(r) & 1-\phi_n(r) \end{bmatrix}.$$

As $P_{1:n+1} = P_{1:n}P_{n:n+1}$, we obtain the following:

$$\begin{aligned} P_{1:n+1} &= \begin{bmatrix} 1-\phi_n(r) & \phi_n(r) \\ \phi_n(r) & 1-\phi_n(r) \end{bmatrix} \begin{bmatrix} 1-r_n & r_n \\ r_n & 1-r_n \end{bmatrix} \\ &= \begin{bmatrix} 1-r_n-\phi_n(r)+2r_n\phi_n(r) & r_n-2r_n\phi_n(r)+\phi_n(r) \\ r_n-2r_n\phi_n(r)+\phi_n(r) & 1-r_n-\phi_n(r)+2r_n\phi_n(r) \end{bmatrix}. \\ &= \begin{bmatrix} 1-\phi_{n+1}(r) & \phi_{n+1}(r) \\ \phi_{n+1}(r) & 1-\phi_{n+1}(r) \end{bmatrix}, \end{aligned}$$

establishing the claim in Equation (21).

The DTMC $J$ satisfies the following property (Kulkarni 2016):

$$\Pr(J_i = j) = \left(\alpha^\top P_{1:i}\right)_j \qquad \forall i \in \{2,3,\dots,N\}, j \in \{0,1\}, \tag{22}$$

where $\alpha^\top = [\alpha_0, \alpha_1]$ is the vector of initial probabilities and $\left(\alpha^\top P_{1:i}\right)_j$ denotes the $(j+1)$-th component of the row vector $\alpha^\top P_{1:i}$. Thus, for every $i \in \{2,3,\dots,N\}$,

$$\begin{bmatrix} \Pr(J_i=0) \\ \Pr(J_i=1) \end{bmatrix}^\top = \begin{bmatrix} \alpha_0 \\ \alpha_1 \end{bmatrix}^\top \begin{bmatrix} 1-\phi_i(r) & \phi_i(r) \\ \phi_i(r) & 1-\phi_i(r) \end{bmatrix} = \begin{bmatrix} \alpha_0+(\alpha_1-\alpha_0)\phi_i(r) \\ \alpha_1+(\alpha_0-\alpha_1)\phi_i(r) \end{bmatrix}^\top.$$

Proposition 1 follows by noting that $\alpha_0 + \alpha_1 = 1$. □

#### Proof of Theorem 1

We use the definition in Equation (14) in the main article to find a closed-form expression for the ECV. Let $L^1$ and $L^2$ be the genotype matrices for the selected pair of individuals, and let $J^1$ ,$J^2$ and $J^3$ be three independent samples from the inheritance distribution. We know that $g^3 = \text{gam}\left(\left[g^1, g^2\right], J^3\right)$ where $g^1 = \text{gam}\left(L^1, J^1\right)$ and $g^2 = \text{gam}\left(L^2, J^2\right)$. Based on the definition of inheritance distribution in Definition 3 in the main article, we have,

$$g_i^1 = L^1_{i,1}(1 - J_i^1) + L^1_{i,2}J_i^1 \qquad \forall i \in [N], \tag{23}$$

$$g_i^2 = L^2_{i,1}(1 - J_i^2) + L^2_{i,2}J_i^2 \qquad \forall i \in [N], \text{ and,} \tag{24}$$

$$g_i^3 = g_i^1(1 - J_i^3) + g_i^2 J_i^3 \qquad \forall i \in [N]. \tag{25}$$

Substitutions in Equation (25) using Equations (23) and (24) yields the expected cross value for the target trait as:

$$\begin{aligned}
\mathbb{E}\left(\sum_{i=1}^{N} g_i^3\right) &= \mathbb{E}\left(\sum_{i=1}^{N} \left[L^1_{i,1}(1 - J_i^1) + L^1_{i,2}J_i^1\right](1 - J_i^3) + \left[L^2_{i,1}(1 - J_i^2) + L^2_{i,2}J_i^2\right] J_i^3\right) \\
&= \mathbb{E}\left(\sum_{i=1}^{N} L^1_{i,1} + \left(L^1_{i,2} - L^1_{i,1}\right)J_i^1 - L^1_{i,1}J_i^3 - \left(L^{k,}_{i,2} - L^1_{i,1}\right)J_i^1 J_i^3 + \right. \\
&\qquad \left. L^2_{i,1}J_i^3 + \left(L^2_{i,2} - L^2_{i,1}\right)J_i^2 J_i^3\right) \\
&= \sum_{i=1}^{N} \left[L^1_{i,1} + \left(L^1_{i,2} - L^1_{i,1}\right)\mathbb{E}(J_i^1) - L^1_{i,1}\mathbb{E}(J_i^3) - \left(L^1_{i,2} - L^1_{i,1}\right)\mathbb{E}(J_i^1 J_i^3) + \right. \\
&\qquad \left. L^2_{i,1}\mathbb{E}(J_i^3) + \left(L^2_{i,2} - L^2_{i,1}\right)\mathbb{E}(J_i^2 J_i^3)\right].
\end{aligned}$$

From Proposition 1 we know that,

$$\mathbb{E}(J_i^1) = \mathbb{E}(J_i^2) = \mathbb{E}(J_i^3) = \alpha_1 + (\alpha_0 - \alpha_1)\phi_i(r) \qquad \forall i \in [N].$$

As $J^1$, $J^2$ and $J^3$ are independent, we know that,

$$\mathbb{E}(J_i^1 J_i^3) = \mathbb{E}(J_i^1)\mathbb{E}(J_i^3) = (\alpha_1 + (\alpha_0 - \alpha_1)\phi_i(r))^2 \qquad \forall i \in [N],$$
$$\mathbb{E}(J_i^2 J_i^3) = \mathbb{E}(J_i^2)\mathbb{E}(J_i^3) = (\alpha_1 + (\alpha_0 - \alpha_1)\phi_i(r))^2 \qquad \forall i \in [N].$$

Thus,

$$\mathbb{E}\left(\sum_{i=1}^{N} g_i^3\right) = \sum_{i=1}^{N}\Big(L_{i,1}^1 + [\alpha_1 + (\alpha_0 - \alpha_1)\phi_i(r)](L_{i,2}^1 - 2L_{i,1}^1 + L_{i,1}^2) + [\alpha_1 + (\alpha_0 - \alpha_1)\phi_i(r)]^2(L_{i,2}^2 + L_{i,1}^1 - L_{i,2}^1 - L_{i,1}^2)\Big). \qquad (26)$$

Assuming $\alpha_0 = \alpha_1 = 0.5$ based on Mendel's second law, Equation (26) reduces to Equation (15) claimed in the main article. □

### FORMULATIONS

#### Mathematical formulation for single-trait parental selection

Following Han et al. (2017), we use the following notations in our integer programming (IP) formulation (27). We use ECV as our objective function and add constraints to restrict inbreeding.

**Parameters:**

- $K \in \mathbb{Z}_{\geq 0}$: Number of individuals in the population
- $N \in \mathbb{Z}_{\geq 0}$: Number of QTL for the target trait
- $G$: $K \times K$ genomic matrix of inbreeding values with elements $g_{k,k'}$ for $k, k' \in [K]$
- $\epsilon \in \mathbb{R}_+$: Inbreeding tolerance on a pair of selected individuals

**Decision variables:**

- $t \in \mathbb{B}^{2\times K}$ representing the parental selection decision where,

$$t_{m,k} = \begin{cases} 1, & \text{if } k\text{-th individual is selected as } m\text{-th parent,} \\ 0, & \text{otherwise,} \end{cases} \quad \forall m \in [2],\ k \in [K].$$

- $x \in \mathbb{B}^{N\times 4}$ representing genotypes of selected individuals. If we suppose that the $k$-th and $k'$-th individuals are selected as first and second parents respectively, i.e., $t_{1,k} = 1$ and $t_{2,k'} = 1$, then:

$$x_{i,j} = L^{k}_{i,j}, \qquad \forall i \in [N],\ j \in \{1,2\},$$
$$x_{i,j} = L^{k'}_{i,j}, \qquad \forall i \in [N],\ j \in \{3,4\}.$$

**Objective function:** Using Equation (15) in the main article, the ECV can be expressed as a function of the decision variables as: $f(t,x) = 0.25 \sum_{i=1}^{N} \sum_{j=1}^{4} x_{i,j}$.

**Formulation:**

$$\max\ 0.25 \sum_{i=1}^{N} \sum_{j=1}^{4} x_{i,j} \tag{27a}$$

$$s.t. \quad \sum_{k=1}^{K} t_{m,k} = 1 \qquad \forall m \in [2] \tag{27b}$$

$$x_{i,j} = \sum_{k=1}^{K} t_{1,k} L_{i,j}^{k} \qquad \forall i \in [N], j \in \{1,2\} \tag{27c}$$

$$x_{i,j} = \sum_{k=1}^{K} t_{2,k} L_{i,j-2}^{k} \qquad \forall i \in [N], j \in \{3,4\} \tag{27d}$$

$$t_{1,k} + t_{2,k'} \leq 1 \qquad \forall k, k' \in [K] \text{ such that } g_{k,k'} \geq \epsilon \tag{27e}$$

$$t_{m,k} \in \{0,1\} \qquad \forall m \in [2], k \in [K] \tag{27f}$$

$$x_{i,j} \in \{0,1\} \qquad \forall i \in [N], j \in [4] \tag{27g}$$

The objective function (27a) maximizes the ECV. Constraint (27b) ensures that exactly two individuals will be selected for the crossing. Constraints (27c) and (27d) assign genotypic information in genotype matrices of the selected individuals to the $x_{i,j}$ variables. Constraint (27e) ensures that two individuals with genomic relationship coefficient greater than the tolerance $\epsilon$ will not be selected simultaneously as parents. As the genomic relationship coefficient between any individual with itself has the highest value of one, this set of constraints will prevent self-crossing between individuals for any value of $\epsilon$ less than one. Finally, constraints (27f) and (27g) force decision variables to take binary values.

#### Algorithm for selecting multiple parental pairs

Suppose we are interested in finding $n_c$ different parental pairs from the population. Assuming that self-crossing is not allowed, we denote the number of feasible solutions (crosses) by $n_f$, which is bounded above by $\binom{K}{2}$. As we impose a constraint for controlling inbreeding, the number of feasible crosses might be strictly less than $\binom{K}{2}$. Specifically, the number of feasible solutions (feasible crosses) is precisely half the number of off-diagonal elements in matrix $G$ that are smaller than $\epsilon$.

If there is no element in matrix $G$ that is smaller than $\epsilon$, then $n_f = 0$ and formulation (27) is infeasible. In this case, we need to increase the value of tolerance $\epsilon$ such that there might be at least $n_f$ possible crosses for the selection. Then, any positive integer value for $n_c$ such that $n_c \leq n_f$ is suitable for our approach.

Assume that after solving the single-trait formulation (27), we find that in the optimal solution we have $t^*_{1,k} = t^*_{2,k'} = 1$. This solution means that $k$-th and $k$'-th individuals are optimal parents that should be crossed. To obtain another pair of parents from the model, we can add the following "conflict constraints" to the single-parent single-trait formulation (27):

$$t_{1,k} + t_{2,k'} \leq 1 \text{ and } t_{1,k'} + t_{2,k} \leq 1. \tag{28}$$

These two constraints will exclude this pair of individuals, $k, k'$, from being selected if we reoptimize formulation (27) with these additional conflict constraints. We can repeat this procedure to find $n_c$ pairs by accumulating the appropriate set of conflict constraints corresponding to individuals selected in the previous iteration. The procedure is summarized in Algorithm 1.

**Algorithm 1** Finding multiple pairs for the parental selection problem

```
1: Input: Appropriate n_c (assumed to be no larger than n_f), G, P, ϵ
2: Output: Set S of selected parental pairs
3: S ← ∅
4: while |S| < n_c do
5:     Solve formulation (27) and obtain optimal solutions t*_{1,k} = t*_{2,k'} = 1.
6:     Add the pair {k, k'} to set S.
7:     Update the formulation by adding the constraints: t_{1,k} + t_{2,k'} ≤ 1, t_{1,k'} + t_{2,k} ≤ 1.
8: end while
9: return S
```

#### Mathematical formulation for multi-trait parental selection

**Additional parameters:**

- $M \in \mathbb{Z}_{\geq 0}$: Number of target traits for the breeding program
- $N_\ell \in \mathbb{Z}_{\geq 0}$: Number of QTL for the $\ell$-th trait $\forall \ell \in [M]$

**Additional decision variables:**

- $x^\ell \in \mathbb{B}^{N_\ell \times 4}$ representing genotypes of selected individuals for each trait $\ell \in [M]$. If we suppose $k$-th and $k'$-th individuals are selected as first and second parents, so $t_{1,k} = 1$ and $t_{2,k'} = 1$, then:

$$
\begin{aligned}
x^\ell_{i,j} &= L^{k,\ell}_{i,j} && \forall i \in [N_\ell], j \in \{1,2\}, \ell \in [M], \\
x^\ell_{i,j} &= L^{k',\ell}_{i,j} && \forall i \in [N_\ell], j \in \{3,4\}, \ell \in [M].
\end{aligned}
$$

**Objective function:** We define the ECV corresponding to the $\ell$-th trait as a function of the decision variables as: $f_\ell(t, x^\ell) = 0.25 \sum_{i=1}^{N_\ell} \sum_{j=1}^{4} x^\ell_{i,j}$. The components of the objective function vector $F(t,x) = \langle f_1(t,x^1), \ldots, f_M(t,x^M) \rangle$ are in decreasing order of importance. Thus, trait $\ell$ is more important than trait $\ell + 1$, for each $\ell \in [M-1]$. Note that we denote the collection of variables $\langle x^1, \ldots, x^M \rangle$ succinctly as $x$.

**Formulation:**

$$\text{lexmax } F(t,x) = \langle f_1(t,x^1),\dots,f_M(t,x^M)\rangle, \tag{29a}$$

$$s.t. \quad \sum_{k=1}^{K} t_{m,k} = 1 \qquad \forall m \in \{1,2\} \tag{29b}$$

$$x_{i,j}^{\ell} = \sum_{k=1}^{K} t_{1,k} L_{i,j}^{k,\ell} \qquad \forall i \in [N_\ell], j \in \{1,2\}, \ell \in [M] \tag{29c}$$

$$x_{i,j}^{\ell} = \sum_{k=1}^{K} t_{2,k} L_{i,j-2}^{k,\ell} \qquad \forall i \in [N_\ell], j \in \{3,4\}, \ell \in [M] \tag{29d}$$

$$t_{1,k} + t_{2,k'} \leq 1 \qquad \forall k, k' \in [K] \text{ such that } g_{k,k'} \geq \epsilon \tag{29e}$$

$$t_{m,k} \in \{0,1\} \qquad \forall m \in \{1,2\}, k \in [K] \tag{29f}$$

$$x_{i,j}^{\ell} \in \{0,1\} \qquad \forall i \in [N_\ell], j \in [4], \ell \in [M] \tag{29g}$$

The multi-objective optimization formulation (29) for the multi-trait parental selection problem lexicographically maximizes the vector of ECV functions corresponding to each trait. We describe this approach in greater detail in the next section. Constraint (29b) states that exactly two individuals will be selected for crossing. Constraints (29c) and (29d) will assign genotypes of selected individuals to $x_{i,j}^{\ell}$ variables. Constraint (29e) implies that any two individuals with an inbreeding coefficient greater than tolerance $\epsilon$ can not be selected as parents for the crossing program. Note that since the inbreeding coefficient between any individual and itself has the highest value (which equals one), for any value of $\epsilon$ less than one, this set of constraints will prevent self-crossing between individuals. Finally, constraints (29f) and (29g) enforce decision variables to take binary values.

#### Lexicographic multi-objective optimization with degradation tolerances

Define a vector of tolerances $\tau = (\tau_1, \tau_2, \ldots, \tau_M)$ such that $\tau_\ell \in [0,1]$ for all $\ell \in [M]$. Since we do not need degradation for the last objective, we set $\tau_M = 0$. Tolerance $\tau_\ell$ represents the allowable degradation for the $\ell$-th objective function. Let us assume that $\chi^1$ is the set of feasible solutions based on the constraints of formulation (29). Let $z_1^*$ be the optimal objective value for the first objective function $f_1(t, x^1)$ over all feasible solutions in set $\chi^1$. That is,

$$z_1^* = \max\{f_1(t, x^1) \mid (t, x) \in \chi^1\}. \tag{30}$$

As the tolerance for the first objective is $\tau_1$, the set of feasible solutions for the second objective is given by:

$$\chi^2 = \{(t, x) \in \chi^1 \mid f_1(t, x^1) \geq (1 - \tau_1) z_1^*\}, \tag{31}$$

and the best objective value for the second objective function is:

$$z_2^* = \max\{f_2(t, x^2) \mid (t, x) \in \chi^2\}. \tag{32}$$

Generally, the set of feasible solutions for the $\ell + 1$-th objective function and its best objective value are as follows:

$$\chi^{\ell+1} = \{(t, x) \in \chi^\ell \mid f_\ell(t, x^\ell) \geq (1 - \tau_\ell) z_\ell^*\} \qquad \forall \ell \in [M-1], \tag{33}$$

$$z_{\ell+1}^* = \max\{f_{\ell+1}(t, x^{\ell+1}) \mid (t, x) \in \chi^{\ell+1}\} \qquad \forall \ell \in [M-1]. \tag{34}$$

The set of "tolerance-optimal" solutions for the problem is given by:

$$\chi^* = \underset{(t,x)\in\chi^M}{\operatorname{argmax}} f_M(t, x^M). \tag{35}$$

By construction, the feasible sets satisfy the following relationship:

$$\chi^* \subseteq \chi^M \subseteq \chi^{M-1} \subseteq \cdots \subseteq \chi^2 \subseteq \chi^1. \tag{36}$$

An optimal solution in multi-objective optimization is called an *efficient* or *non-dominated* solution based on some domination structure chosen by the decision maker (Sawaragi et al. 1985). Arguably, the most well-known notion of efficiency is Pareto optimality. We say that the solution $(\hat{t}, \hat{x})$ is Pareto optimal (or non-dominated) if there is no feasible solution $(t, x)$ to Formulation (29) such that $F(t, x) \geq F(\hat{t}, \hat{x})$ and $f_\ell(t, x) > f_\ell(\hat{t}, \hat{x})$ for at least one trait $\ell$ (Miettinen et al. 2016).

As we show using the next result, the set of tolerance-optimal solutions $\chi^*$ is guaranteed to contain Pareto optimal solutions. Furthermore, if $(t, x) \in \chi^*$ then, either $(t, x)$ is Pareto optimal or it is dominated by a Pareto optimal solution in $\chi^*$.

**Proposition 1.** *Every solution $(t^*, x^*) \in \chi^*$ is either Pareto optimal, or it is dominated by a Pareto optimal solution in $\chi^*$.*

*Proof.* If $(t^*, x^*) \in \chi^*$ is not Pareto optimal, then there exists a Pareto optimal solution $(t', x') \in \chi^1$ that dominates $(t^*, x^*)$. Note that the existence of such a solution follows from the finiteness of $\chi^1$. We prove that $(t', x') \in \chi^*$ by contradiction.

Suppose $(t', x') \notin \chi^*$. Then, there exists $i \in [M]$ such that $(t', x') \in \chi^i$ and $(t', x') \notin \chi^{i+1}$ (where $\chi^{M+1} = \chi^*$). Therefore,

$$f_i(t', x'^i) < (1 - \tau_i) z_i^*. \tag{37}$$

If $i = M$, we arrive at a contradiction because inequality (37) implies that $(t', x')$ does not dominate $(t^*, x^*)$. (Recall that $\tau_M = 0$.)

Now suppose, $i \in [M-1]$. By Equation (36), we know that $(t^*, x^*) \in \chi^* \subseteq \chi^{i+1}$, and that,

$$(1-\tau_i)z_i^* \leq f_i(t^*, x^{*i}). \tag{38}$$

Inequalities (37) and (38) imply that,

$$f_i(t', x'^i) < f_i(t^*, x^{*i}). \tag{39}$$

Again, contradicting the assumption that $(t', x')$ dominates $(t^*, x^*)$. This implies that *every* Pareto optimal solution $(t', x')$ that dominates $(t^*, x^*)$ belongs to $\chi^*$. □

Based on Proposition 1, if we seek a Pareto optimal solution, we can guarantee the identification of one by carrying out an additional step and solving one more optimization problem given by:

$$\max \left\{ \sum_{\ell \in [M]} f_\ell(t, x^\ell) \mid (t, x) \in \chi^* \right\}.$$
